# The role of transposon activity in shaping cis-regulatory element evolution after whole genome duplication

**DOI:** 10.1101/2024.01.02.573861

**Authors:** Øystein Monsen, Lars Grønvold, Alex Datsomor, Thomas Harvey, James Kijas, Alexander Suh, Torgeir R. Hvidsten, Simen Rød Sandve

**Author notes:** Present address of Alexander Suh: Centre for Molecular Biodiversity Research, Leibniz Institute for the Analysis of Biodiversity Change, Adenauerallee 160, D-53113 Bonn, Germany. contributed equally (Øystein Monsen, Lars Grønvold).

## Abstract

Two of the most potent drivers of genome evolution in eukaryotes are whole genome duplications (WGD) and transposable element (TE) activity. These two mutational forces can also play synergistic roles; WGDs result in both cellular stress and functional redundancy, which would allow TEs to escape host-silencing mechanisms and effectively spread with reduced impact on fitness. As TEs can function as, or evolve into, TE-derived cis-regulatory elements (TE-CREs), bursts of TE-activity following WGD are likely to impact evolution of gene regulation. However, the role of TEs in genome regulatory remodelling after WGDs is unclear. Here we used the genome of Atlantic salmon, which is known to have experienced massive expansion of TEs after a WGD ∼100 Mya, as a model system to explore the synergistic roles of TEs and WGDs on genome regulatory evolution.

We identified 55,080 putative TE-CREs in Atlantic salmon using chromatin accessibility data from brain and liver. Of these, 80% were tissue specific to liver (43%) or brain (37%) and TE-CREs originating from retroelements were twice as common as those originating from DNA elements. Signatures of selection shaping TE-CRE evolution were evident from depletion of TEs in open chromatin, a bias in tissue-shared TE-CREs towards older TE-insertions, as well as tissue-specific processes shaping the TE-CRE repertoire. A minority of TE-families (16%) accounted for the origin of 46% of all TE-CREs, but the transposition activity of these ‘CRE-superspreader’ families happened mostly prior to the WGD. Analyses of individual TE-CREs do however support a significantly higher rate of TE-CRE evolution from insertions happening around the time of the salmonid WGD. This pattern was particularly striking for the DTT elements, despite having generally low propensity to evolve into TE-CREs and impact transcription. Furthermore, co-expression based analyses supported the presence of TE-driven gene regulatory network evolution, including DTT elements active at the time of WGD.

In conclusion, we find a strong association between TE insertions at the time of WGD and TE-CRE evolution. This association was not driven by particular TE-families with high capability to evolve into TE-CREs but likely a consequence of the concurrent surge of novel TE insertions, mostly from DTT elements, in combination with a shift in selective pressure on genome regulation following the WGD.

## Introduction

The two most influential mutational mechanisms that have shaped eukaryotic genome evolution are whole genome duplications (WGD) and transposable element (TE) activity. Both WGDs and TEs drive genome size evolution. However, as mobile genetic elements with capacity to replicate (Feschotte and Pritham 2007), TEs also impact genome evolution in numerous other ways, by generating novel genes (Elisaphenko et al. 2008; Qin et al. 2015; Diehl et al. 2020; Cosby et al. 2021), modulating chromatin looping (Diehl et al. 2020), rearranging genome structure (Bourque et al. 2018) as well as supplying “raw material” for gene regulatory evolution in the form of cis-regulatory elements (CREs) (Bourque et al. 2008; Feschotte 2008; Sundaram et al. 2014; Chuong et al. 2017; Cosby et al. 2019; Diehl et al. 2020; Sundaram and Wysocka 2020).

Studies of mammalian genomes have provided deep insights into the role of TEs in CRE-evolution and the potency of TE-derived CREs (TE-CREs) to regulate gene expression (reviewed in Fueyo et al. (2022)). For example, as much as 40% of the mouse and human transcription factor (TF) binding sites have been shown to be within TEs (Sundaram et al. 2014), and as many as 19% of pluripotency factor TFs are located within TEs (Kunarso et al. 2010; Sundaram et al. 2017). Curiously, in mammals TEs associated with gene regulation during development have been shown to be younger than those associated with regulation in adult somatic tissues (reviewed in (Fueyo et al. 2022)), suggesting different evolutionary pressures on TEs with distinct regulatory roles.

Genome evolution through TE activity is also likely influenced by WGDs. Because WGDs result in cellular stress, TEs can escape host-silencing mechanisms following WGDs. This is supported by both experimental (Kashkush et al. 2002, 2003; Kraitshtein et al. 2010) and comparative genomics (Lien et al. 2016; Marburger et al. 2018) studies. Additionally, WGDs result in increased functional redundancy. This will reduce the average negative fitness effects of novel TE insertions and thereby allow for fixation of TE insertions following WGD (Baduel et al. 2019), including insertions that influence gene regulation. In line with this, Gillard et al. (Gillard et al. 2021) recently reported that TE insertions in promoters were associated with regulatory divergence of gene duplicates following WGD in salmonid fish. However, systematic investigations into the role of TEs in CRE evolution and genome regulatory remodelling after WGDs are still lacking.

Here we address this knowledge gap regarding the role of WGD in TE-associated genome regulatory evolution using salmonids as a model system. Salmonids underwent a WGD 80-100 Mya (Lien et al. 2016) which coincided with the onset of a burst of TE activity, particularly featuring elements belonging to the DTT/Tc1-mariner superfamily. This observation has led to the hypothesis that increased TE activity in the immediate aftermath of the WGD was a major driver of genome regulatory evolution. To explore this idea we leverage ATAC-seq data from two tissues (brain and liver) to identify putative CREs that have evolved from TE-derived sequences. We then combine these TE-CRE annotations with analyses of the temporal dynamics of TE activity, analyses of gene-coexpression, and massive parallel reporter assays. Our results support a link between WGD and TE-CRE evolution, and support the idea of synergistic interactions between WGDs and TE activity to drive rewiring of genome regulation.

## Results

### The TE-CRE landscape of Atlantic salmon

To investigate the contributions of different TEs to CRE evolution, we first characterised the TE landscape of the salmon genome using an updated version of the existing TE annotation from (Lien et al. 2016). The total TE-annotation covered 51.92% of the genome. Consistent with previous findings (Goodier and Davidson 1994; Lien et al. 2016), the dominating TE group was DNA transposons from the Tc1-Mariner superfamily with >655,000 copies, covering 327 million base pairs, just shy of 10% of the genome (Figure 1A-C). In general, the genomic context of TE insertions was quite similar to the genomic baseline (Figure 1D), but with slightly more TEs in intronic regions and slightly less TEs in exons and intergenic regions. Of the well-represented TE superfamilies (>10k insertions) only the Nimb retrotransposon superfamily was an exception to this pattern, for which 18% of the copies were found in promoter regions (Figure 1D).

**Figure 1.**
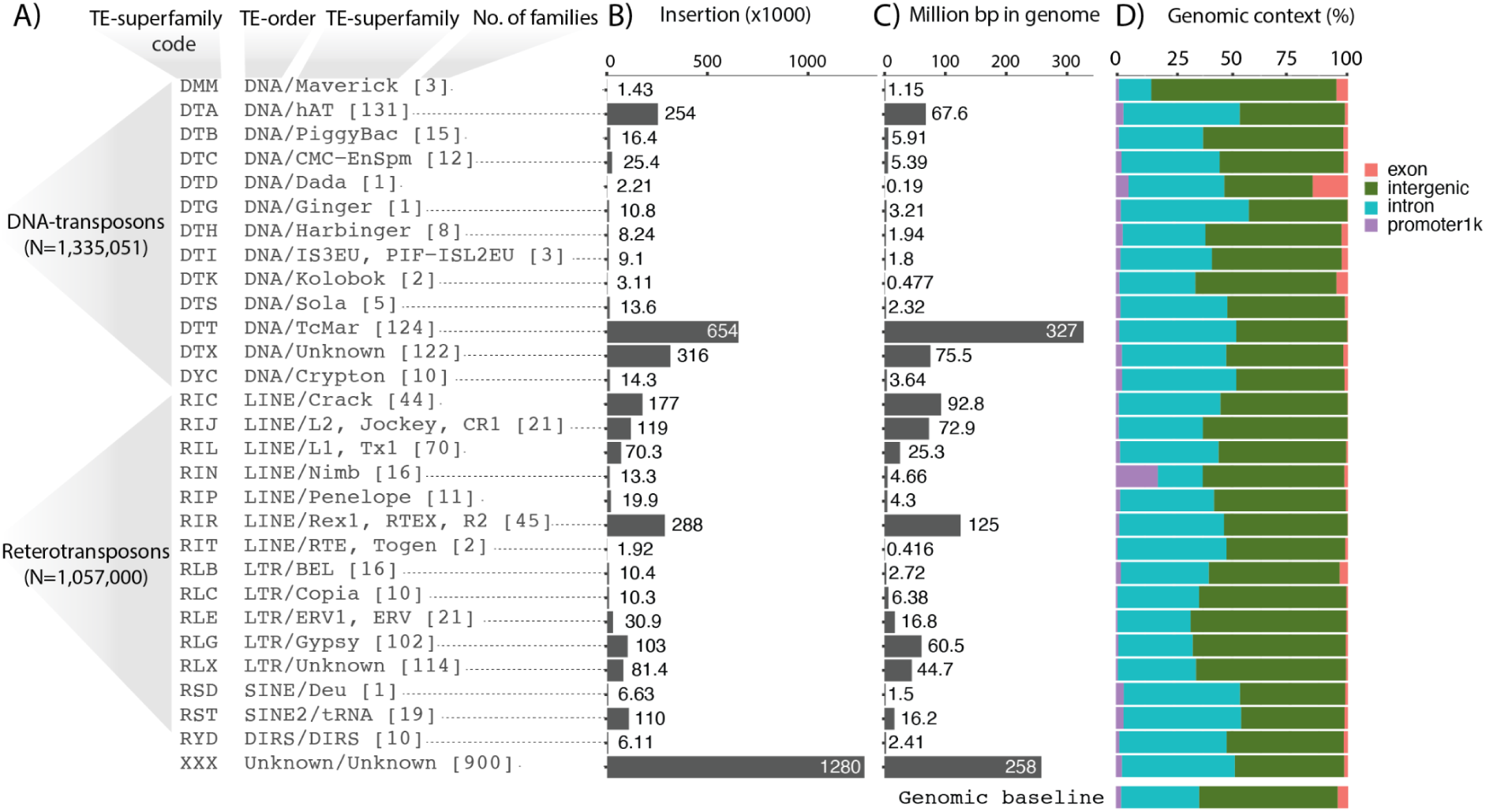
Overview of the genomic TE landscape. **A)** Superfamily level overview of TE annotations in the Atlantic salmon genome. Number of TE-families per superfamily in square brackets. **B)** TE insertions per superfamily. **C)** Annotated base pairs at the TE superfamily level. **D)** TE annotations (bp proportions) overlapping different genomic contexts. Genomic baseline is the proportion of the entire genomic sequence that is assigned to the four genomic contexts.

Active CREs in tissues and cells are associated with increased chromatin accessibility (Keene et al. 1981; McGhee et al. 1981; Buenrostro et al. 2013). Thus, to study the contribution of TEs to the salmon CRE landscape, we integrated our TE annotation with annotations of accessible chromatin regions identified using ATAC-seq data from liver and brain. Analysis of the overlap between TEs and accessible chromatin revealed a large depletion of TEs in accessible chromatin. While TEs represent ∼52% of the genome sequence, TE insertions only represent <20% of the regions of accessible chromatin (Figure 2A), with liver having a higher proportion of annotated TEs in accessible chromatin than brain.

**Figure 2.**
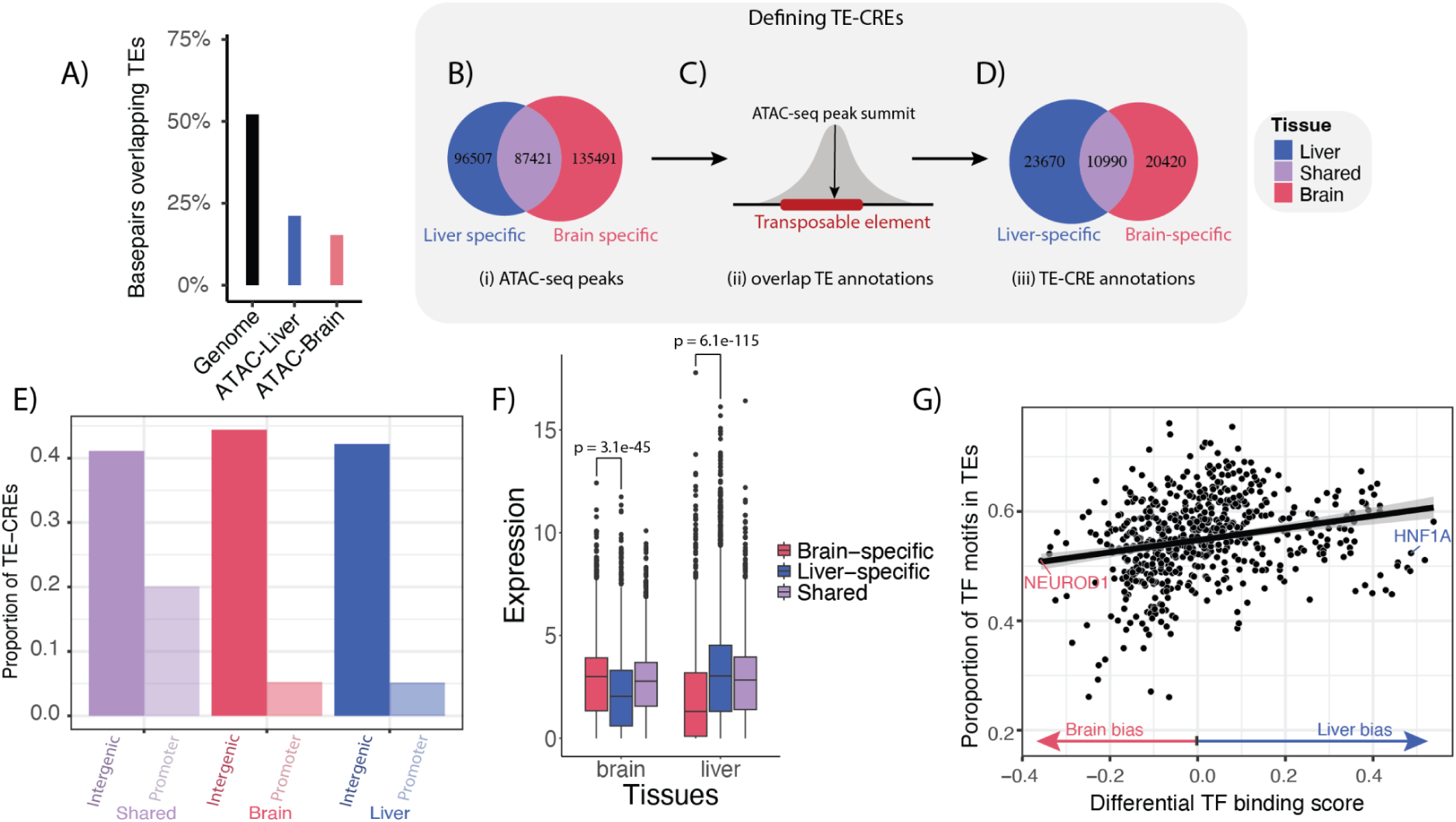
TE-CRE landscape. A) The proportion of base pairs overlapping TEs, either out of all genome-wide bp or those within an ATAC-seq peak. B-D) Pipeline to define putative TE-CREs. B) Venn diagram of tissue-specific and shared ATAC-peaks from liver and brain. C) Cartoon showing how TE-CREs are defined as ATAC-seq peak summits when overlapping with a TE. D) Venn diagram of tissue-specific and shared TE-CREs from liver and brain. E) Proportion of shared and tissue-specific TE-CREs in promoter vs. intergenic regions. F) Gene expression levels of the nearest genes to tissue-specific and shared TE-CREs in brain and liver. P-values from Wilcoxon-test indicated above tissues. G) Correlation between TF tissue specificity and the proportion of genome wide TF motif matches located in TEs. Each point represents a TF motif. Tissue specificity is based on differential TF binding score from TOBIAS (Bentsen et al. 2020), which essentially summarises the relative ATAC-seq footprint signal across all potential binding sites.

To define a set of TEs that contribute to putative CREs, we narrowed in on those TE annotations overlapping chromatin accessibility peaks (Figure 2B-D). These were defined as putative TE-CREs. Although the majority (55%) of TE annotations (excluding ‘unknown’ repeats without classification) were DNA elements (1,335,051 insertions), TE-CREs from DNA elements were a minority (27%). Both the proportion of basepairs in accessible chromatin (Figure 2A) and number of putative TE-CREs (Figure 2D) were higher in the liver compared to the brain. Of a total of 55,080 TE-CREs, 20% were shared between tissues, 37% were brain-specific, and 43% were liver-specific (Figure 2D). Tissue-shared TE-CREs were overrepresented about 4-fold in promoters compared to the tissue-specific TE-CREs (Figure 2E). We also found that tissue-specific TE-CREs were associated with tissue-bias in gene expression (Figure 2F), supporting a regulatory effect of TE-CREs.

One reason for TE-CREs tending to be specific to liver rather than brain (Figure 2D) could be due to differences in purifying selection pressure on regulatory networks important for brain function compared to liver function. One expectation from this hypothesis would be that TF binding motifs with strong brain bias in TF binding would be depleted in TE sequences. To test this, we first inferred tissue bias in TF binding using genome-wide TF binding motif occupancy signals through TF-footprinting. We then correlated these signals with the proportion of TF binding motifs found in TE sequences. In line with our expectations, motifs for brain biassed TFs were less frequently found in TEs (regression line in Figure 2G). The most highly liver-biassed TFs, such as HNF1A, were exceptions to this general trend, although these liver-biassed TFs were fewer and much less depleted in TEs compared to the most highly brain-biassed TFs such as NEUROD1 (Figure 2G). Taken together, our results are in line with a model whereby TEs with smaller chances of having a negative fitness consequence for the host are evolutionarily more successful.

### A minority of TEs have CRE superspreader abilities

Next we wanted to understand the contribution of specific TE-families to the TE-CRE landscape. Overall, there was a positive linear relationship between the genomic copy number and the number of TE-CRE for TE-superfamilies (Figure 3A). However, some superfamilies (Figure 3A, see data points outside 95% CI), contributed significantly less (RIC and DTT) or more (RIN, RLG, DTA, RIJ) to the TE-CRE landscape than expected based on the genomic copy numbers (Figure 3A). Particularly striking was the DTT superfamily elements, which are dominating in terms of numbers of insertions (∼27% of all TE copies with an assigned taxonomy) but only represented ∼4% of the TE-CREs.

**Figure 3.**
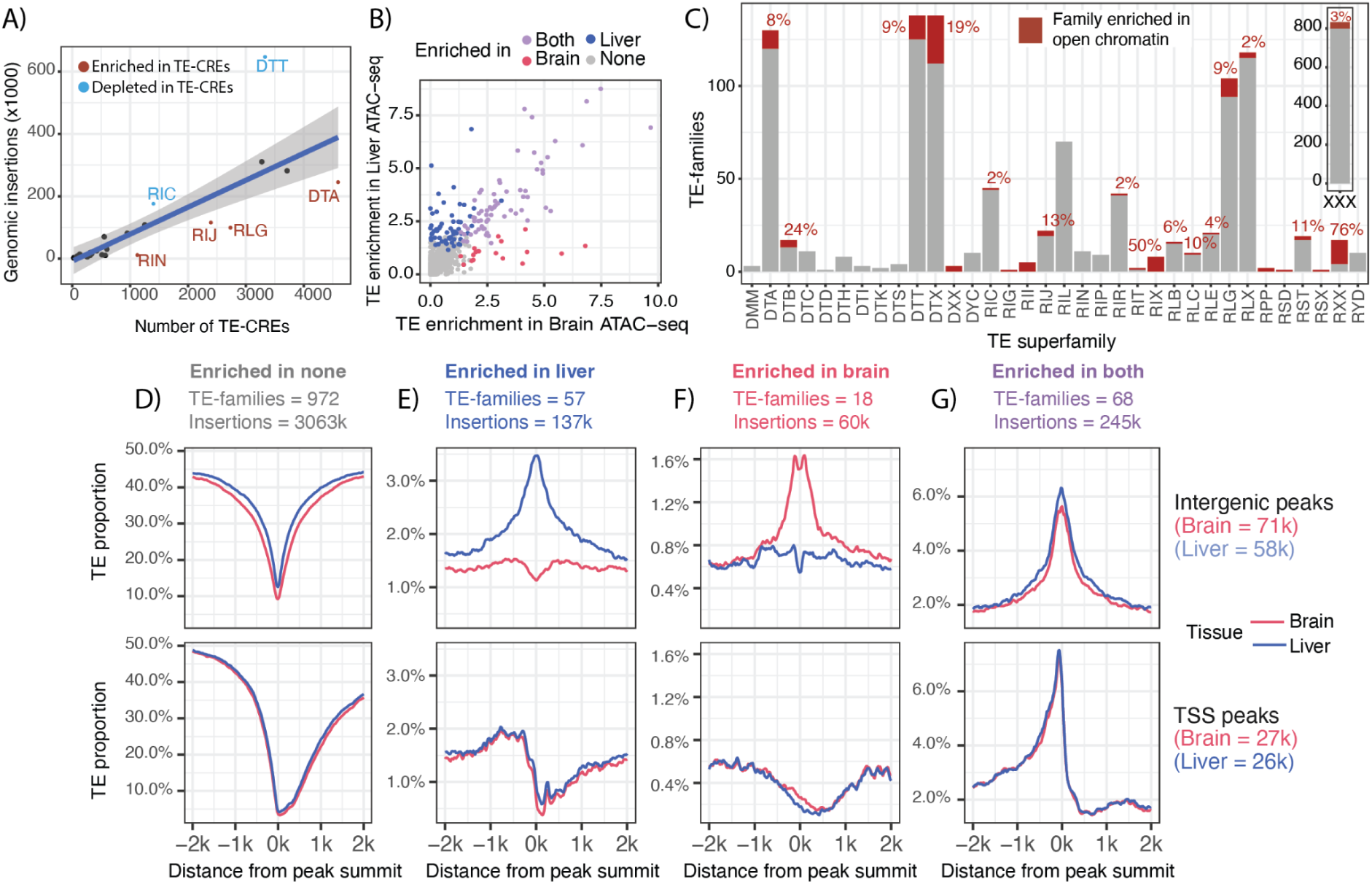
TEs enriched in open chromatin. A) The number of insertions per superfamily plotted against the number of CREs in each superfamily. The shaded area is a 95% confidence level interval. Superfamilies falling outside the 95% confidence interval is annotated with the three letter superfamily code B) TE-families (>500 genomic insertions) plotted according to fold-enrichment within ATAC-seq peaks in brain and liver. TE families are assigned into categories based on enrichment in liver, brain or both. C) Proportion of TE-families enriched in open chromatin per superfamily. A manual curation step of the TE-families enriched in open chromatin resulted in a slightly different superfamily list than the initial machine-predicted annotations presented in Figure 1. Note also that only TE-family sequences with > 500 insertions have been included. The percent enriched TE-families per superfamily are indicated above bars. D-G) Proportion of bp overlapping TEs from each enrichment category around peak summits in intergenic or promoter regions (summit within 500 bases of TSS). Peaks in promoter regions are oriented according to the corresponding TSS with gene bodies to the right in figures.

To characterise the TE-CRE landscape in more detail, we identified individual TE-families enriched in open chromatin (see methods and Figure 3B). TE-families significantly enriched in open chromatin are hereafter referred to as ‘CRE-superspreaders’. After removing TE-families with few genomic insertions (500 or fewer) we found that only 178 (16%) out of 1119 TE-families were defined as CRE-superspreaders (Figure 3B). Forty nine percent of these superspreaders were enriched in open chromatin in both tissues (88), while 39% (69) and 12% (21) were tissue-specific and enriched in accessible chromatin only in the liver or brain, respectively. The proportion of taxonomically unclassified TE-families (three-letter code “XXX”) was >50% among the identified CRE-superspreaders (101 out of 178 families). For these particularly interesting TE-families we therefore performed manual curation, resulting in four TE-families being discarded from further analyses, and only 34 families remained taxonomically unclassified (Supplementary Table 1).

We find that CRE-superspreaders were taxonomically diverse, belonging to 18 different TE superfamilies, but that very few TE-families in the DTT superfamily evolved into CRE-superspreaders (Figure 3C). Note that superfamilies consisting only of CRE-superspreader TEs is a technical artefact stemming from the manual curation of the taxonomically unknown (three-letter code XXX) superspreaders.

Next we explored the local TE landscape around the open chromatin peaks in various genomic contexts (intergenic and promoters) and tissues (Figure 3 D-G). We find that in promoters the proportion of TEs in open chromatin decreases towards TSS and the gene body, probably reflecting increased purifying selection pressure (less tolerance for TE-insertions) inside the gene body.

Furthermore we find that close to genes (i.e. in promoters), TE proportions were higher for tissue-shared (Figure 3G) compared to tissue-specific (Figure 3E-F) TE-CREs. In intergenic regions (i.e. putative enhancers) we find very strong tissue-specific TE enrichment signals not present in promoter TE-CREs (Figure 3 E-F). In sum, we find that CRE-superspreader TEs are biassed towards certain taxonomic groups of TEs and that these TEs are enriched in accessible chromatin with distinct patterns and effect sizes across tissues and genomic contexts.

### The temporal dynamics of TE-CRE evolution

The main hypothesis we set out to test in this study was whether the increase in TE-activity associated with salmonid WGD was instrumental in driving TE-CRE evolution. To explore this hypothesis we first calculated sequence divergence between individual TE-insertions and their TE-family consensus sequence in the form of CpG-normalised Kimura distances. Sequence divergence estimates can be used as a proxy for time, and then be compared to the Kimura distance expectation for TEs being active prior to or after the salmonid WGD event (see methods for details). One challenge with sequence divergence based comparisons is however the intrinsic connection between sequence divergence and purifying selection pressure that can bias our results. Here we used the entire TE insertion (not only the part that is in open chromatin) to estimate divergence to TE-family consensus, hence we expect this bias to be negligible. Nevertheless, we first analysed the sequence divergence distributions of different classes of TE-CREs as well as TE sequences not in accessible chromatin (Figure 4A). As expected, we do not find that TE-CREs are more similar to their TE-family consensus than other TE insertions, supporting that putative purifying selection on TE-CRE function does not transfer to our sequence divergence based age-proxy. If anything, the TE sequences giving rise to tissue-shared CREs are older than TEs not giving rise to TE-CREs (see ‘both’ in Figure 4A).

**Figure 4.**
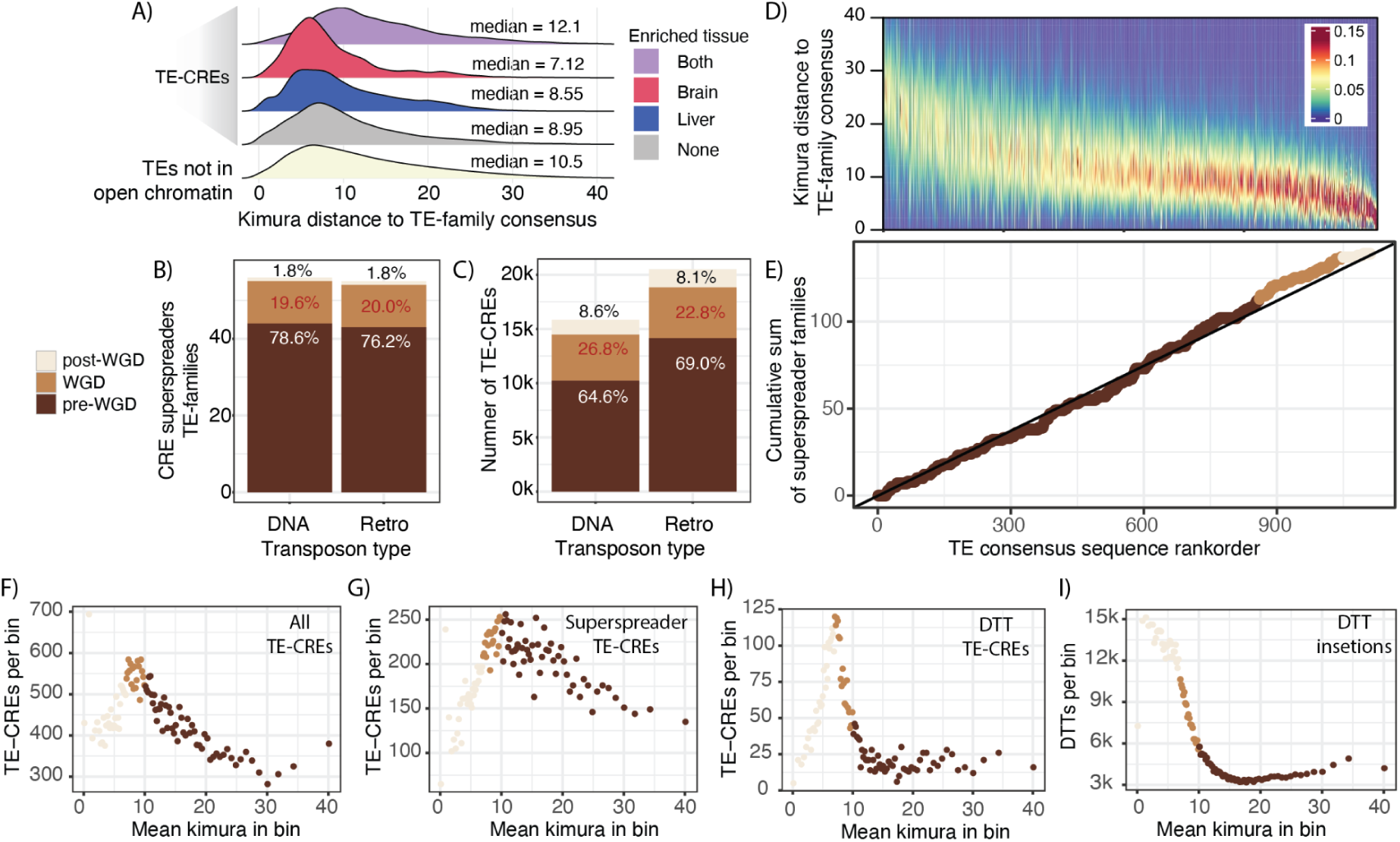
Temporal dynamics of TE-CRE insertion activity by TE taxonomy. A) Distribution of sequence divergence of TE-CREs from their TE-family consensus sequence. Colours represent if TE-CRE are from TE-families with superspreader ability (liver, brain, or both) or not (grey). B) Number of TE-families with superspreader ability subdivided into DNA- and retroelements. Colours represent the TE-family age proxy calculated as mean divergence between genomic insertions and their consensus TE sequence. Post-WGD = <7 Kimura distance, WGD = 7-10 Kimura distance, pre-WGD = >10 Kimura distance. C) Number of TE-CREs from TEs with a taxonomic classification subdivided into DNA- and retroelements. Colours represent the TE-family age proxy. D) Heatmap of the divergence distributions of all insertions per TE-family (with >500 insertions) to their consensus sequence. TE-families are ordered based on mean divergence from consensus. E) Cumulative distribution of CRE-superspreader TE-families ordered by meanKimura distance between genomic copies and TE-family consensus sequence. Colours represent age proxy as defined by mean divergence to TE-family consensus sequence F-I) The number of TE-CREs (F-H) and TE insertions (I) per ‘age’-bin of Kimura distances for all TE-CREs, TE-CREs from superspreader families, and TE-CREs from the DTT superfamily.

Next, we stratified the TE-CREs on their transposition age-proxy relative to the WGD and their taxonomic order (Figures 4B and 4C). Seventeen percent (189) of TE-families had a mean Kimura distance between insertions and the TE-family consensus reflecting activity in the approximate range around the time of the WGD (Kimura distance of 7-10). Among the TE-families with CRE-superspreader ability, there were about the same proportion of TE-families active around the time of WGD classified as DNA-elements (19.6%) compared to retroelements (20.0%) (Figure 4B). Slightly higher proportion of TE-CRE insertions from DNA-elements were assigned to the time around WGD based on the mean Kimura distance to TE-family consensus (Figure 4C, 26.8% vs 22.8%), however the retroelement-derived TE-CREs dominated over DNA-element TE-CREs in absolute numbers (Figure 4C).

Even if most TE-CREs are older than the salmonid WGD, it is still plausible that the WGD triggered a shift in the evolutionary rates of TE-CREs. To explore this in more detail we first analysed the temporal dynamics of all TE-families (>500 genomic copies, Figure 4D) by plotting the cumulative sum of TE-CRE superspreader families against all TE-families ordered by mean divergence to consensus (Figure 4E). If WGD were associated with a general burst of CRE-superspreader activity we expect to see a steeper slope in the cumulative sum distribution around mean kimura distances of 7-10 to the TE-family consensus. Although we find a slight change in CRE-superspreader accumulation rates around this divergence range (Figure 4E), most of the data points lie on or close to the diagonal (null model) and the age distribution of CRE-superspreaders TE-families was not significantly different from other TE-families (two sided Kolmogorov-Smornov test, p-value = 0.5). Nor did we find a significant increase in the proportion of TE-CRE superspreader families among TE-families with predicted activity around the WGD (mean kimura distance 7-10, Fisher test, p-value = 0.45). Taken together, these results do not support a model whereby the WGD caused a large shift in TE-CRE superspreader family activity.

Our family-level analyses showed that CRE-superspreaders activity were evenly distributed in time and mostly pre-WGD (Figure 4B, E). Yet, the distributions of Kimura distances for TE-CRE insertions (except those from families enriched in both tissues) suggests that TE-CREs are biassed towards insertions happening around the time of, or more recent than the WGD (median Kimura distance <10, Figure 4A). One explanation for this seeming contradiction could have been that younger TE-families have more insertions that evolve into TE-CREs, but this is not the case (Figure 4C). Another explanation could be that using the mean of each TE-family obscures within family heterogeneity in Kimura distances which can arise from multiple activity bursts or prolonged activity across time. As an alternative approach to associate WGD with TE-CRE evolution we therefore used the Kimura divergence from individual TE insertions. We divided TE insertions into 100 equally large ‘age’ bins based on Kimura distance and plotted the distribution of TE-CREs per bin. For all TE-CREs we found a significant enrichment (Fisher exact test, p=3.4e100) of TE-CREs with a Kimura distance to consensus reflecting activity around the time of the WGD (Figure 4F). Using only TE-CREs from superspreader TE-families, a much weaker association with the WGD was found (Figure 4G), with more TE-CREs coming from TE activity prior to the WGD (in line with Figure 4B). Tallying TE-CREs per superfamily into our three defined temporal activity periods (Supplementary figure 2) we find that the DTT superfamily has the largest proportion of TE-CREs arising from insertions at the time of the WGD. In fact, the Kimura distances of DTT-derived TE-CREs (Figure 4H) reflects an extremely strong clustering of TE-CREs with insertion ages close to the WGD. This pattern could be caused by a simple correlation between the number of DTT insertions and number of TE-CREs, but this is not the case as the total number of DTT insertions continues to increase as Kimura distances decrease towards zero, while TE-CRE numbers drop dramatically for DTT insertions after the WGD (Figure 4I). In conclusion, we find that the WGD is strongly associated with increased rates of TE-CRE evolution, particularly from DTT elements, but that this association is not driven by higher transposition activities of CRE-superspreader TE-families.

### Co-expression analysis support TE-CRE driven regulatory network evolution

If TEs are spreading CREs with sequences that either have a potent TF binding motif or are prone to mutate into a TF motif, we expect different genes with similar TE-CREs (TEs insertions belonging to the same consensus sequence) to be more similarly regulated than random gene pairs. To identify such putative cases of TE-CRE driven evolution of gene regulation, we assigned each TE-CRE to the closest gene and tested if genes with similar TE-CREs were more co-expressed than expected by chance.

We first used RNA-seq data from the liver of 112 individuals spanning different ages, sex and different diets in fresh water (Gillard et al. 2018). In the context of this liver co-expression network, significant co-expression (low p-values) indicate that TE-CREs from one particular TE-family are candidates for modulating the gene regulation in the liver depending on developmental and physiological states. Using only TE-CREs from liver, 41 TE-families (41 of 1387 = 3%) were associated with genes that were significantly co-expressed (FDR-corrected p-value < 0.05) (Figure 5A). Of these significant TE-families, 20 (49%) were CRE superspreaders. The significant TE-families came from 12 TE superfamilies and TEs of unknown origin (XXX) accounting for 29% (Figure 5B). The cumulative distribution of TE-CREs associated with gene co-expression did not suggest a temporal co-occurence of WGD and the TE-CREs with putative gene regulatory effects (Figure 5C).

**Figure 5.**
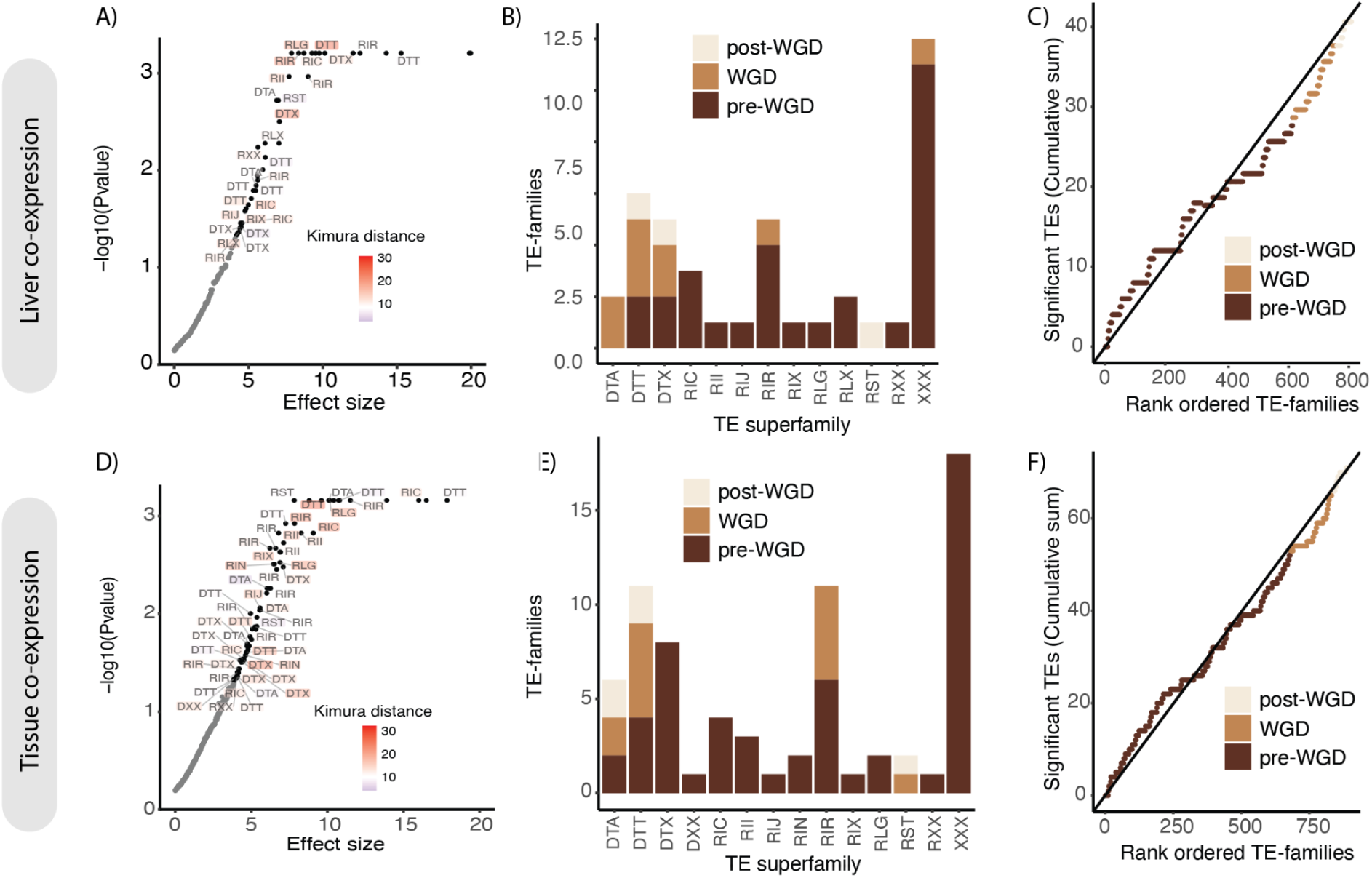
TE-CREs driving co-expression. Top row A-C shows results from liver co-expression. Bottom rows D-F shows results from tissue atlas co-expression. A and D) Significance (FDR-adjusted p-values) plotted against effect size (standard deviations) for each TE-family, indicating the strength of co-expression of their associated genes in the liver (B) and tissue atlas (D) co-expression networks, respectively. Points with fdr-adjusted p-value < 0.05 are labelled with superfamily names (unknown/XXX superfamilies are not labelled). B,E) Distribution of significant TE-families grouped by superfamilies in liver (B) and tissue atlas (E) data sets. C and F) Cumulative distribution of TE-families with significant effect on gene co-expression in liver (C) and tissue atlas (F) data sets. Temporal classification was based on the mean divergence of all TE insertions to their TE consensus sequence where post-WGD was defined as Kimura distance < 7, WGD as 7-10 and pre-WGD as >10.

TE-CREs are also known to induce tissue-specific regulatory effects (Karttunen et al. 2023). We therefore conducted the same analyses using RNA-seq data from 13 different tissues (Lien et al. 2016). Using TE-CREs from both the liver and brain, 71 TE-families (71 of 1465 = 4.8%) were associated with significant co-expression (Figure 5D), of which 29 (41%) were superspreaders. The significant TE-families came from 13 TE superfamilies (Figure 5E). Each significant TE-family was associated with a tissue TE-CRE-profile (fraction of TE-CREs found in liver, brain or both), and these profiles generally agreed with the tissue expression profiles of the associated genes (RNA-seq expression values across 13 tissues), thus corroborating that our approach indeed identified regulatory-active TE-CREs. Similar to the liver co-expression analyses, the cumulative distribution of TE-CREs impacting tissue-regulation did not suggest any link between the WGD and the TE-CRE evolution shaping gene co-expression (Figure 5F). Taken together, we find evidence for a small proportion (3-5%) of the TE-families spreading CREs that regulate nearby genes.

### Functional validation of TE-CREs using massively parallel reporter assay

To be able to directly assess regulatory potential of TE-CREs in Atlantic salmon we carried out massive parallel reporter assays (MPRA), specifically an ATAC-STARR-seq experiment, in salmon primary liver cells (Figure 6A). This method assesses the ability of random DNA fragments from accessible chromatin to modulate transcription levels (Wang et al. 2018). In total, 4,267,201 million unique DNA fragments from open chromatin in liver were assayed. Thirty four percent of these fragments (1,456,914) could be assigned to a specific TE insertion site (>50% overlap with a TE annotation) (Figure 6B). Of the TE-derived sequence fragments assayed, 1.2% had transcriptional regulatory activity, a slightly lower proportion than non-TE fragments (1.6%) and, TE-derived regulatory active fragments were more likely to induce transcription compared to non-TE sequences (see “Up” in Figure 6C).

**Figure 6.**
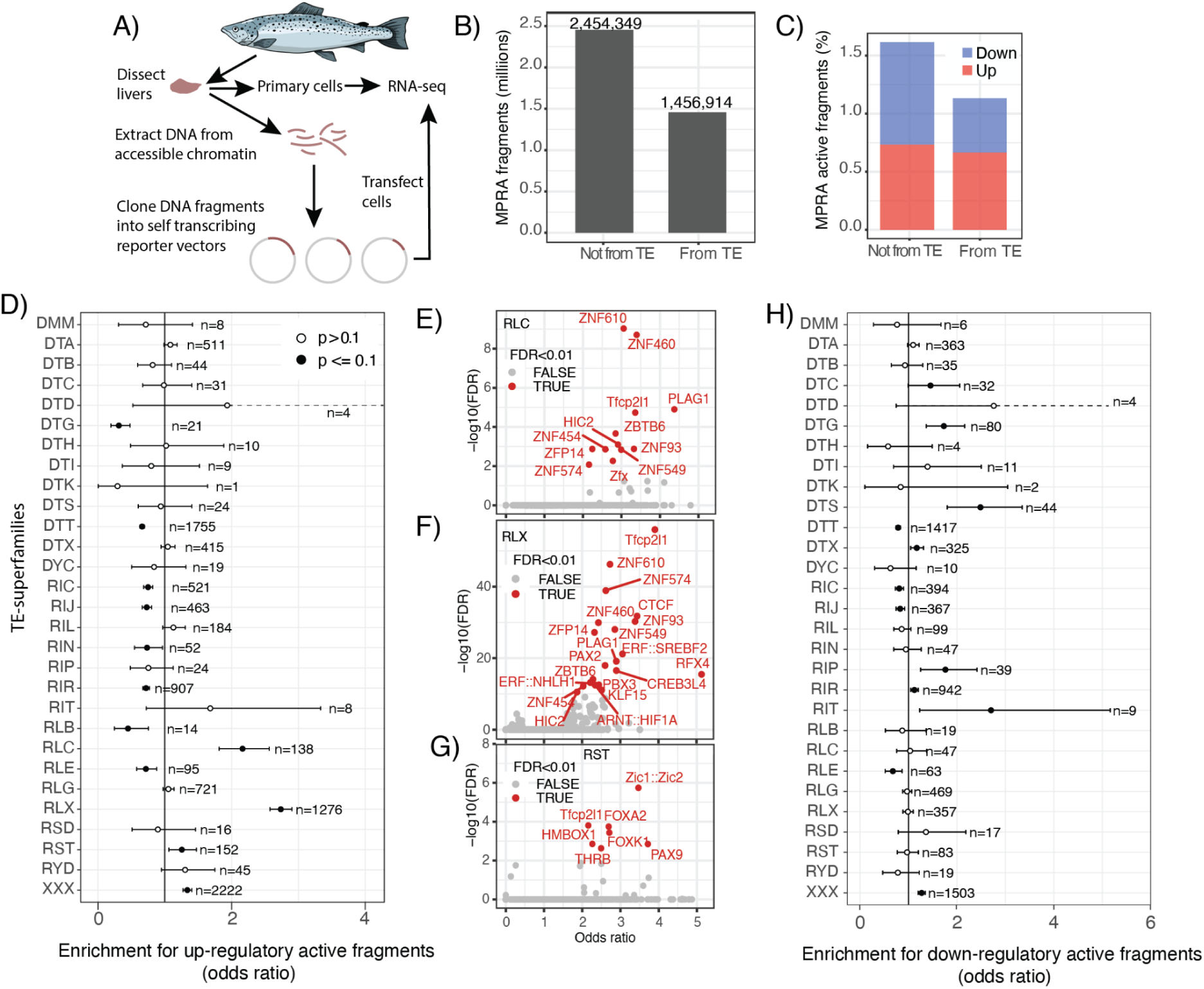
Massive parallel reporter assay screening of regulatory activity. A) schematic overview of the ATAC-STAR-seq MPRA experiment. B) Barplot of the origin of sequence fragments included in the analyses. C) Regulatory activity (inducer or repressor) of MPRA sequence fragments from TE and non-TE sequences. D) Fisher test results for enrichment of transcriptional inducing MPRA fragments within a TE-superfamily compared to all other TEs. Unknown taxonomy and DNA/retrotransposons of unknown origin (DTX/RLX) are considered separate groups. Number of regulatory active fragments are given for each category (n). E-G) TF motif enrichment in transcriptionally inducing MPRA fragments from TE superfamilies enriched in regulatory active fragments. TF names are from the JASPAR database and the nomenclature reflects whether it came from human or mouse. H) Fisher test results for enrichment of transcriptional repressing MPRA fragments within a TE-superfamily compared to all other TEs. Unknown taxonomy and DNA/retrotransposons of unknown origin (DTX/RLX) are considered separate groups. Number of regulatory active fragments are given for each category (n).

To test if CRE-sequences from particular TE insertions were more likely to increase gene expression (i.e. act as enhancers) we compared the ratio of regulatorily active vs inactive fragments. This analysis was done only at the TE-superfamily level (including groups with partially assigned and unknown taxonomy) since there were too few unique fragments from each TE-family. These analyses revealed clear differences between superfamilies (Figure 6D). Three retrotransposon superfamilies were significantly enriched for regulatorily active fragments (Fisher test, fdr-corrected p-value < 0.05). Two of these were LTRs (RLC and RLX) which had >2-fold higher ratio of fragments acting as enhancers, while another SINE superfamily (RST) was significantly enriched but with a much lower effect size estimate (Figure 6D). The transcriptionally inducing fragments from these three superfamilies were enriched for a total of 38 unique TF binding motifs (RLC=12, RST=8, and RLX = 19) (Figure 6E-G). Many of these top-enriched motifs belong to known liver active TFs (i.e. Srebf2REBF2, KlfLF15, FoxaOXA2, ThrbHRB) (Tao et al. 2013; Lau et al. 2018; Chaves et al. 2021; Yerra and Drosatos 2023), including the tfcp2l1 motif (Wei et al. 2019) which were enriched in all three superfamilies (Figure 6E-G). We also tested for enrichment of transcriptionally repressing activity and found 6 superfamilies (in addition to the XXX and DTX groups) that were enriched for transcription repressing fragments (Figure 6H).

Interestingly, fragments from the WGD-associated DTTs were significantly less likely to be regulatorily active (both to induce and repress transcription) compared to other TE-derived sequences in the MPRA experiment (i.e. odds ratio < 1 in Figures 6 D and H). This is consistent with our findings that DTTs are depleted in TE-CREs compared to random expectations (Figure 3A).

## Discussion

### The Atlantic salmon TE-CRE landscape

Most in-depth characterizations of TE-associated CREs have so far been carried out in mammalian cells and tissues. Our investigations into the Atlantic salmon genome revealed similarities with mammals, but also highlighted some unique features of the salmonid TE-CRE landscape. About ∼15-20% of CREs were derived from TE-sequences (Figure 2A, 2C), which is in the lower bound of what has been found in mammals using similar methods to identify TE-CREs (Bourque et al. 2008; Kunarso et al. 2010; Sundaram et al. 2014). Consistent with studies of mammalian genomes (Simonti et al. 2017; Nishihara 2019), the majority of putative TE-CREs in Atlantic salmon were associated with enhancer function rather than promoters (Figure 2E).

Mammalian TE repertoire (Feschotte and Pritham 2007) and TE-CRE landscapes (Nishihara 2019; Pehrsson et al. 2019; Roller et al. 2021) are dominated by retroelements. In most fish (Shao et al. 2019), including Atlantic salmon (Figure 1, DNA transposons = 55% of the TEs), DNA transposons are the dominating TEs. However, similar to mammals the majority of Atlantic salmon TE-CREs (73%/45,419) were derived from retroelements (Figure 4C). Our MPRA data (Figure 6) also pointed to retroelements being more likely to induce transcription compared to DNA transposons (Figure 6 D) and that transcription-inducing fragments from these TEs were enriched for TF binding motifs known to be bound by liver-active TFs (Tao et al. 2013; Lau et al. 2018; Chaves et al. 2021; Yerra and Drosatos 2023). Only one TF binding motif, the tfcp2l1, was enriched across all three superfamilies enriched for transcription-inducing fragments (Figure 6 E-G). Tfcp2l1 has previously been found to bind LTRs in human stem cells (Wang et al. 2014) and is proposed to be a top regulator of human hepatocyte differentiation (Wei et al. 2019). Hence the tfcp2l1 stands out as a key player in shaping evolution of retroelement-associated TE-CRE landscapes in Atlantic salmon.

Although retroelements dominate the salmon TE-CRE landscape, the role of DNA transposons such as DTA and DTT elements in TE-CRE evolution cannot be neglected due to their high genomic copy numbers (Figure 1, Figure 3A). Indeed, the TE superfamily contributing the highest number of TE-CREs was the DTA (hATs) superfamily of DNA elements (Figure 3A). DTAs have also been found important for TE-CRE evolution in several other species. Enrichment of DTA element-insertions in accessible chromatin has also been found in maize (Noshay et al. 2021), and DTA elements make up a significant proportion (15%) of the TE-derived CTCF sites associated with TAD loop anchoring in certain human cell types (Choudhary et al. 2023). In this study we also find families of DTA (as well as DTT TEs) driving rewiring of tissue gene regulatory networks (Figure 5B, E). Furthermore, even though DTA sequences were not significantly more likely to drive transcription compared to any other TE superfamily (fdr-corrected p-value = 0.18, Figure 6D), DTA sequences were more likely to induce transcription (0.72% of fragments were up-regulatory active) compared to sequence fragments derived from DNA transposons in general (0.51% up-regulatory active). Hence, DNA transposons have been a considerable source of novel CREs sequences and likely played an important role in the evolution of genome regulation in Atlantic salmon and other salmonids.

### Selection on TE-CRE repertoire

Studies examining how evolutionary forces mould the genomic TE-landscape underscore the significant role of purifying selection in limiting TE accumulation within protein-coding gene sequences. (Bartolomé et al. 2002; Rizzon et al. 2003), but also in non-coding regions (Hollister and Gaut 2009; Bergthorsson et al. 2020; Langmüller et al. 2023). These selection signatures on TE insertions in non-coding regions indicate selective forces on TE-CRE evolution, which is also evident from several analyses in our study.

We find clear underrepresentation of TE sequences in accessible chromatin (Figure 2A), and in particular near the peaks in accessible chromatin in promoters and intergenic regions (Figure 3D), consistent with purifying selection against TE accumulation in regulatory active regions (Bergthorsson et al. 2020; Langmüller et al. 2023).

In mammals, TEs-CRE are typically from older TE insertions (Simonti et al. 2017; Pehrsson et al. 2019) suggesting that selection pressure on TEs depend on TE insertion age, which is likely related to deterioration of transposition ability as TEs age and accumulate mutations. In Atlantic salmon however, we do not find a general trend of older TE-sequences giving rise to TE-CREs (Figure 4A). This could be linked to a general relaxation of purifying selection pressure after WGD (Ronfort 1999; Baduel et al. 2019), see section below for in depth discussion. However, we do find that tissue-shared TE-CREs clearly have an older origin compared to tissue-specific TE-CREs (Figure 4A). One way to interpret this age bias is that tissue-specific TE-CREs have on average more neutral fitness effects. Conversely, older and tissue-shared TE-CREs are more likely to be advantageous, fixed by selection, and maintained for longer under purifying selection. Under this model we expect higher TE-CRE turnaround rates (loss and gain) for tissue-specific compared to tissue-shared TE-CREs, which has been described in mammals (Roller et al. 2021). Higher evolutionary turnaround rates of tissue-specific TE-CREs is also expected if tissue- or cell-type specific CREs are ‘easier’ to evolve than tissue-shared CREs, which has recently been suggested to be the case (Luthra et al. 2022).

Since gene regulation is under tissue-specific selection pressure (Brawand et al. 2011; Berthelot et al. 2018), we expect CRE-evolution to be under different selection pressures in different tissues. From mammalian studies we know that purifying selection on gene regulation is stronger in the brain than liver (Wang et al. 2020), hence we expect TE-CRE evolution to reflect this asymmetry in selection pressure. Consistent with this expectation we find clear tissue differences in TE-CRE numbers (Figure 2D) and that TE sequences were consistently depleted in highly brain biassed TF binding motif (Figure 2G). One interpretation of these results may be that the evolutionary arms race between genomic ‘parasites’ and the host results in selection pressure to “avoid” having sequences that function as, or can evolve into CREs that impact gene regulatory networks related to critical brain-functions. Since the liver is a key organ for nutrient conversion and detoxification, it is also possible that higher rates of liver biassed TE-CRE evolution (compared to brain) reflects adaptive evolution of liver function as a response to continuous changes in the environment through macroevolutionary timescales.

### TE-CRE evolution in aftermath of the WGD

The whole genome duplication in the ancestor of salmonids resulted in large scale gene regulatory rewiring (Lien et al. 2016; Varadharajan et al. 2018). These novel gene regulatory phenotypes have been partly linked to divergent TE-insertions in promoters of gene duplicates (Gillard et al. 2021; Sahlström et al. 2023), but the link between WGD and TE-CRE evolution has remained elusive. One hypothesis is that WGD induce a genomic shock which results in bursts of TE activity (the ‘genomic shock’ model (McClintock 1984)), and that these novel TE insertions allow for rapid TE-CRE evolution and rewiring of gene regulatory networks in the initial aftermath of a WGD. Another hypothesis is that relaxed purifying selection in polyploids allows for higher rates of TE accumulation (Baduel et al. 2019), which in turn will lead to higher rates of neutral and nearly-neutral TE-CRE evolution. In this scenario, however, there is no expectation of a temporal link between bursts of TE-activity and bursts of TE-CRE evolution.

Our results do lend support to the ‘genomic shock’ model for TE-CRE evolution following WGD, as we find an increase in the rate of TE-CRE evolution from insertions happening around the time of WGD (Figure 4F). It is however important to note that the WGD-associated TE-CRE evolution is not driven by specific TE-families that were particularly effective at evolving into TE-CREs (Figure 4B, C, G). DTTs, which exploded in numbers around the WGD (Lien et al. 2016), were poor at evolving into TE-CREs (Figure 2G) and significantly less likely to impact transcription compared to other TEs (Figure 6D, H), in line with other studies showing that the DTT superfamily does not contain many TF binding sites (Simonti et al. 2017; Zeng et al. 2018). Despite this, the WGD-associated rate shift in the evolution of TE-CREs (Figure 4H, I) was particularly strong for DTTs, and we also demonstrated DTT-associated rewiring of gene regulatory networks after the WGD (Figure 5). These results could be explained by a transient period of extreme relaxation of selective constraints, or a confined period of strong selection to recalibrate optimal gene expression phenotypes in the aftermath of the genome doubling. However, to further quantify the importance of selection on TE-CRE evolution, a larger comparative approach (Andrews et al. 2023) is warranted.

## Methods

### TE annotation

The TE library (ssal_repeats_v5.1) used to annotate TEs in this study is described in detail in (Richard Minkley 2018). To generate a TE annotation of the salmon genome (ICSASG v2 assembly) we used RepeatMasker version 4.1.2-p1 (Smit et al. 2015) under default settings with the ssal_repeats_v5.1 library. RepeatMasker takes a library of TE consensus sequences and detects whole and fragmented parts of these consensuses across the genome using a BLAST-like algorithm. The output file contains the genomic coordinates of the annotation, and various quality measures such as completeness, and divergence from consensus. The latter measure was used to estimate relative ages of TE activity. TE superfamilies were assigned a three letter tag based on the classifications from Figure 1 in (Wicker et al. 2007). Where there was no obvious categorisation, a literature review was conducted to determine the taxonomic status of a superfamily, and a new tag name introduced based on available letters (so e.g. Nimb is here called RIN as a superfamily of LINE elements).

Manual curation of specific TE families was done following an adapted version of Goubert et al’s process (Goubert et al. 2022), under inspiration from Suh (Suh et al. 2018): Using BLASTn (Altschul et al. 1990), we aligned each TE-consensus to the genome, extracted the twenty best matches and extended them by 2000bp upstream and downstream. We checked the extended matches against the RepBase (Bao et al. 2015) database using BLASTn and xBLAST with standard settings, before we aligned them using MAFFT’s ‘einsi’ variant (Katoh and Standley 2013). Then, we inspected these alignments for structural features in BioEdit (Hall 1999) and, if conservation across the sequence was deemed interesting, in JalView (Waterhouse et al. 2009). In addition, we ran the TE-Aid package (https://github.com/clemgoub/TE-Aid) on each consensus to help guide curation efforts and check each consensus according to its annotation profile and self-alignment. This helped screen for technical noise such as microsatellite sequences near sites of local annotation enrichment. If the annotating consensus was deemed to be incomplete (i.e. if parts of the extended sequence aligned well outside of the consensus), we used Advanced Consensus Generator (https://www.hiv.lanl.gov/content/sequence/CONSENSUS/AdvCon.html) to generate a new consensus from the most complete of the extracted alignments for classification.

We produced plots using base R’s (R Core Team 2021) plot function, as well as the packages ggplot2 (Wickham 2016) and cowplot. Both the Tidyverse (Wickham et al. 2019) and data.table packages were used for analysis, summary statistics and data wrangling.

### Defining genomic context

Based on the NCBI gene annotation (refseq ID: GCF_000233375.1), each part of the genome was assigned as promoter, exon, intron or intergenic. For Figure 1D the promoter was defined as 1000 bp upstream to 200 bp downstream of each transcription start site (TSS). Gene annotations can overlap, e.g. because of multiple transcript isoforms, so overlapping annotations were merged by prioritising promoter > exon > intron > intergenic. For TE-CREs (Figure 2E and 3D-G) each peak was classified as promoter if the summit is less than 500bp upstream or downstream from start of gene (i.e. first TSS per gene) or intergenic if summit is more than 500bp from any gene (exon and intron TE-CREs are not specifically mentioned).

### ATAC-seq peak calling

To annotate regions of accessible chromatin we used ATAC-seq data from four brains and livers from Atlantic salmon (ENA project number PRJEB38052). The ATAC-seq reads were mapped to the salmon genome assembly (ICSASG v2, refseq ID: GCF_000233375.1) using BWA-MEM. Genrich v.06 (https://github.com/jsh58/Genrich) was then used to call open chromatin regions (also referred to as ‘peaks’) with default parameters, apart from ‘-m 20 -j’ (minimum mapping quality 20; ATAC-Seq mode). Genrich uses all four replicates to generate peaks, resulting in one set of peaks for each tissue. The summit of each peak is identified as the midpoint of the peak interval with highest significance.

### TE-CRE definition

To define TE-CREs we combined the ATAC-seq peak set with our TE annotations and classified an ATAC-seq peak as a TE-CRE if the peak summit is inside a TE-annotation. TE-CREs were defined as shared between tissues if (i) the brain ATAC-seq peak summit was within the liver ATAC-seq peak interval and (ii) both the liver and brain peak summits are inside the same TE annotation.

### TF motif scanning

To identify potential TF binding sites in the genome we scanned the entire genome for motif matches using FIMO (Grant et al. 2011). The TF motifs were downloaded from the 2022 JASPAR CORE vertebrates non-redundant motif database (Castro-Mondragon et al. 2022).

### Differential TF binding score between liver and brain

To identify TFs that are specific to liver or brain we used the differential binding score that was generated by the TOBIAS software (Bentsen et al. 2020) in an earlier study (Gillard et al. 2021)the same ATAC-seq data. In short, the differential binding score quantifies the change in TF binding activity between the two tissues by comparing footprints in accessible chromatin regions. TOBIAS first identifies TF footprints based on ATAC-seq signal, representing areas protected by bound TFs. It then performs motif matching against the underlying genomic sequence to associate footprints with specific transcription factors. The differential binding score is then summarised across individual binding sites to provide a global measure of TF activity. In our case, positive scores indicate higher TF binding in liver, while negative scores indicate higher binding in brain.

### Identification of TE families enriched in open chromatin

To identify TE families which had contributed more to TE-CREs than expected by chance we counted the observed number of ATAC-seq peak summits that are inside an annotated TE for each family and compared that to the expected count if the summits were randomly distributed along the genome. However, since some TE families tend to have certain preferences to where they are inserted in the genome (Chuong et al. 2017), we take into account the genomic context when calculating the expected counts following the methodology described in (Bogdan et al. 2019). Specifically, the genome is divided into different regions based on distance from the annotated genes. The regions are (in order of priority): 5UTR, exon, tss (less than 1 kb upstream), promoter (1–5 kb upstream), intron, proximal (5–10 kb upstream or less than 10 kb downstream), distal (10–100 kb upstream or downstream) and desert (greater than 100 kb upstream or downstream). Within each genomic context the expected count was determined by multiplying the number of summits with the proportion of bases covered by the TE family. Note that, because the summits are single base pair points, this straightforward calculation is equivalent to shuffling the summits like they did in (Bogdan et al. 2019). After determining the context specific expected count, the total expected count was calculated as the sum of those. We then used a one-sided binomial test to determine if the observed overlap was significantly greater than expected. Note that the sum of binomial distributions is not exactly a binomial distribution but with large enough numbers, as in our case, the difference is insignificant. The test was performed for each TE-family with more than 500 insertions and p-values were corrected for multiple testing using the Benjamini-Hochberg method. This procedure was repeated for both liver and brain.

### Estimating the temporal activity of TEs

TE-insertions will accumulate individual substitutions and diverge in sequence similarity over time. Hence, the sequence divergence between insertions and their TE-family consensus sequence can be used as a proxy for the time since transposition activity. Sequence divergence estimates were extracted from RepeatMasker software output (Smit et al. 2015). Note that we did not use the default divergence measure reported by RepeatMasker, but rather Kimura distances normalised for CpG content. This adjustment accounts for the high mutation rates at CpG loci due to methylation and heterogeneity in GC content among different TE-families. Specifically, two transition mutations at a CpG site are counted as a single transition, one transition is counted as 1/10 of a standard transition, and transversions are counted as usual. These CpG-adjusted Kimura values can be found in the “.align” files output by RepeatMasker when running with the “-a” option.

Since our TE activity ‘age’ estimates in the form of Kimura distances cannot easily be converted to absolute time, we used an empirically driven approach to define a Kimura distance interval which represented the approximate time of WGD. All TE insertions happening prior to WGD would become represented on two duplicated chromosomes after the WGD. We therefore reasoned that TE-families with a high proportion of their TE-copies in genomic alignments between duplicated regions of the salmon genome would likely represent TE activity from before the WGD event. Conversely, transposition events occurring after the WGD would likely not end up in the paralogous genomic region in the genome, and thus not be included in genomic alignments between duplicated chromosomes. Finally, we plotted the mean Kimura distance between the TE-family consensus sequences and all genomic insertions for that family against the proportion of TE insertions in duplicate region alignments (Supplemental Figure 1). This plot indicated that the proportion of TE insertions in alignments started increasing at a Kimura distance around 7-10, which we then used to have a rough estimate of the time at or just after the salmonid WGD.

### Co-expression analysis

We used two RNA-seq expression data sets to analyse the effect of TE-CREs on gene expression: (1) A liver data set comprising 112 samples spanning different diets and life stages in fresh-water (Gillard et al. 2018) (Gillard et al. 2018)27,786 expressed genes(Gillard et al. 2018) and (2) a tissue atlas comprising 13 different tissues (Lien et al. 2016) (24,650 expressed genes).

TE-CREs in liver, brain or both (the ATAC-seq peak summits of the liver and brain TE-CREs reciprocally overlapped the peak in the other tissue) were assigned to genes with the closest transcription start site (TSS) (no distance threshold was enforced). For each TE-family with insertions associated with at least five expressed genes, we computed the network density of the associated genes (i.e. the mean Pearson correlation between all gene pairs associated with that TE-family). False Discovery Rate (FDR)-corrected p-values were obtained by comparing these network densities to those of randomly selected expressed genes. We ran 100 000 simulations drawing the same number of genes, containing the same number of WGD-derived duplicates (which are often co-expressed), as found in the original data. Effect sizes were calculated as the number of standard deviations away from the mean of randomised network densities.

### Massive parallel reporter assay

Transcriptional regulatory potential of TE-CREs in Atlantic salmon was assessed using ATAC-STARR-seq as previously described in Wang et al. (2018). We used the pSTARR-seq reporter plasmid with the core promoter of Atlantic salmon elongation factor 1 alpha, EF1α (NC_027326.1: 7785458-7785702) instead of the super core promoter 1 (SCP1) originally adapted in human cells (Arnold et al. 2013). ATAC DNA fragments were extracted from Atlantic salmon liver cell nuclei following the OmniATAC protocol (Corces et al. 2017). A clean-up step was performed using Qiagen MinElute PCR purification kit and PCR-amplified using NEBNext Ultra Q5 DNA polymerase master mix (New England Biolabs®) with forward primer (5’-TAGAGCATGCACC GGCAAGCAGAAGACGGCATACGAGAT[N10]ATGTCTCGTGGGCTCGGAGATGT-3’, where N10 corresponds to a random 10 nucleotide i7 barcode sequence) and reverse primer (Rv:5’-GGCCGAATTCGTCGATCGTCGGCAGCGTCAGATGTG-3’). Thermo cycling conditions were 72 ^0^C for 5 min, 98 ^0^C for 30 sec, 8 cycles of 98 ^0^C for 10 sec, 63 ^0^C for 30 sec and 72 ^0^C for 1 min. PCR products were purified using Qiagen MinElute PCR purification kit and size-selected (∼30-280 bp) using Ampure XP beads (Beckman Coulter). Reporter plasmid libraries were made by cloning amplified ATAC fragments into AgeI-HF- and SalI-HF-linearized pSTARR-seq plasmid using InFusion HD cloning kit (Takara) and then propagated in MegaX DH10B T1R electrocompetent bacteria. Plasmids were isolated using the NucleoBond® PC 2000 Mega kit (MACHEREY-NAGEL). An aliquot of plasmid library was PCR-amplified with i5 and i7 primers and sequenced on Novaseq (150 bp Paired-end) and aligned to salmon genome to ensure sufficient complexity and proportions of cloned fragments within open chromatin region. Plasmid library was electroporated into primary salmon hepatocytes as previously described (Datsomor et al. 2022). Total RNA was isolated 24 hours post-transfection using the Qiagen RNeasy Midi columns. Poly A+ RNA from total RNA was extracted using the mRNA isolation kit (Roche). Remaining genomic DNA in isolated mRNA were digested with Turbo DNase (Thermo Fisher). Complementary DNA (cDNA) from mRNA was generated using the Superscript III Reverse transcriptase (Thermo Fisher) with a gene-specific primer (5’-CAAACTCATCAATGTATCTTATCATG-3′). Sequencing-ready libraries from cDNA and the input (reporter plasmid library) were prepared as previously described by Wang et al. (2018) and Tewhey et al. (Tewhey et al. 2016).

Sequenced reads were mapped to the salmon genome assembly (ICSASG v2, refseq ID: GCF_000233375.1) using BWA-MEM. The number of read-pairs mapped to each unique location was counted. Each unique location, i.e. having a specific start and end, was assumed to come from a unique fragment. These counts were fed into DESeq2 using the DNA (input plasmid library) as control and contrasted with the RNA (cDNA) samples. Fragments with significant RNA to DNA ratio were used to define fragments with significant regulatory activity. Prior to DESeq2 the fragment counts were split into bins by length.

### TF motif enrichment in TE-derived active MPRA fragments

To investigate what TF motifs that drive the enhancer activity in TE-CREs we performed a fisher exact test to test the dependence between having a motif in the MPRA fragment and the fragment being regulatory active. For each TE superfamily (RLX, RLC or RST), we considered the MPRA fragments that overlap a TE insertion (with at least 5 base pairs), and checked if it had any TF motif matches in the overlapping region. The fisher exact test was performed for each motif, testing the dependency between the fragment being classified as up-regulating and it having a motif match. False discovery rate was controlled with the Benjamini & Hochberg method.

### Data access

All scripts to reproduce figures and analysis are available at https://gitlab.com/sandve-lab/TE-CRE with a frozen version of the scripts deposited to Zenodo (https://doi.org/10.5281/zenodo.13907938). Supplementary data was also deposited to Zenodo (https://doi.org/10.5281/zenodo.13903583). Raw sequencing data from the ATAC-STARR-seq MPRA experiment has been deposited to the European Nucleotide Archive (https://www.ebi.ac.uk/ena/) under accession number PRJEB81135.

## Acknowledgements and funding

This research was funded by NMBU and the Norwegian Research Council through the projects Transpose (275310), Rewired (274669), and DigiSal (248792). We thank Sigbjørn Lien for comments on earlier versions of the manuscript.

## Author contributions

SRS and TRH conceived the study. SRS and TRH acquired funding. AD and TH performed lab experiments related to the massive parallel reporter assays. ØM, LG, SRS, and TRH performed analyses. ØM, LG, TRH, and SRS drafted the manuscript. All authors took part in critical discussions of various aspects of lab-work and/or analytical approaches relevant to their expertise. All authors critically reviewed the manuscript.

## Supplementary material

Table S1: Curation notes and classification. Every CRE-superspreader TE-consensus has been inspected manually as per the procedure in Materials and Methods. ‘consensus_TE’ is the ID of the annotating consensus in question, ‘original_annotation’ is the automatic classification, ‘manual_curation_three_letter_code’ is the post-curation three-letter ID.

