## Supplementary Figures for "The role of transposon activity in shaping cis-regulatory element evolution after whole genome duplication"

Supplemental figures for Monsen et al.

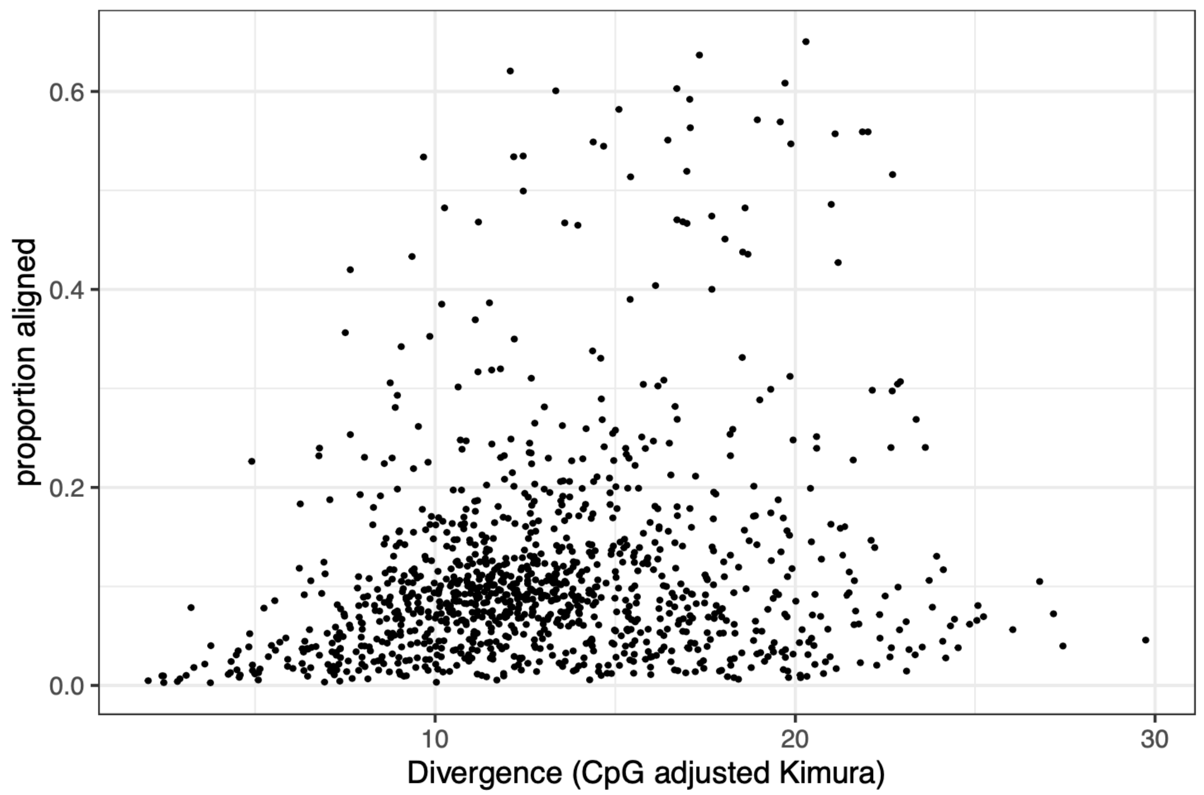


**Supplemental Figure 1. TE family age estimate vs proportion aligned between duplicate regions.** Y-axis shows Mean kimura distance from consensus for each TE family and X-axis shows the proportion of aligned bases between duplicate regions in all TEs from each family.

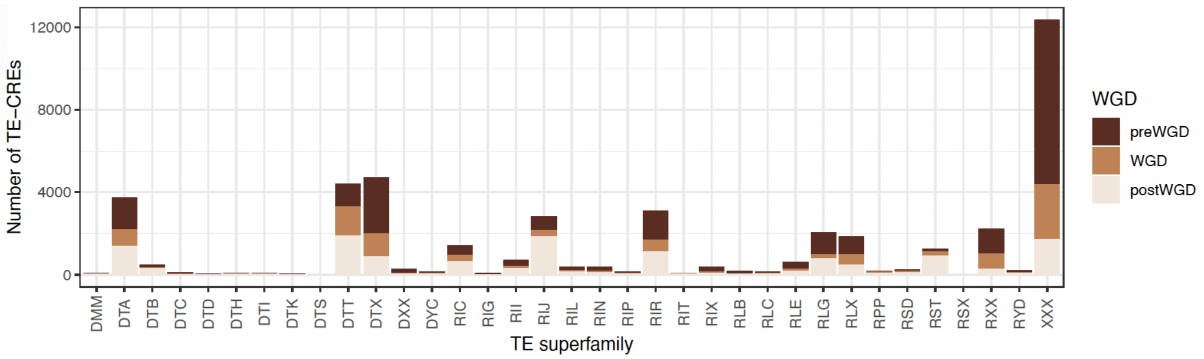


**Supplemental Figure 2. TE-CREs per superfamily.** Colours reflect Kimura distances between each TE-insertion and its TE-family consensus sequences where post-WGD = <7 Kimura distance, WGD = 7-10 Kimura distance, pre-WGD = >10 Kimura distance.
